# A metadata managed FAIR end-to-end workflow for microbial community Omics data analysis

**DOI:** 10.64898/2025.12.22.696032

**Authors:** C. Ke, J.J. Koehorst, B. Nijsse, W.T. Scott, P.J. Schaap

## Abstract

**Background:** Molecular profiling using high-throughput ‘omics technologies has tremendously increased our ability to interrogate complex microbial communities at the molecular level. In the context of data reuse, the FAIRification of these extensive datasets is frequently perceived as a secondary administrative task, addressed only after data analysis has been completed. However, this approach overlooks the potential benefits of early metadata integration as the procedures for processing and analyzing raw data are primarily dictated by the underlying research design and experimental conditions. Gathering interoperable research metadata at the earliest stages creates a standardized basis for managing, processing, and analyzing data enabling more efficient and reproducible FAIR workflows.

**Results:** The single containment principle was used to develop modular containerized reproducible workflows that support the FAIR principles for research software by systematically capturing standardized metadata for each data processing step along with the resulting data products. Using defined mock metagenomic datasets as an example, we show that interoperable research metadata can be used to drive such computational workflows. By processing raw data accordingly, machine-actionable provenance chains are created that enhance the reproducibility and reusability of the resulting data products.

**Conclusions:** A seamless integration of wet lab experiments with computational investigations is essential for a FAIR end-to-end research process. Meta-data-managed workflows prevent the need for unnecessary data manipulation. Workflow provenance registration explicates the complex multi-step methods employed for data processing and analysis. Combining FAIR principles with data provenance registration enhances the reusability of omics datasets by promoting transparency and reproducibility.

Key Points

- We present the first metadata-managed FAIR end-to-end workflow for microbial community Omics data analysis.
- The framework links experiment metadata with CWL workflows, enabling seamless metadata-driven execution from experiment to data product.
- Comprehensive provenance tracking using RDF standards creates machine-actionable chains connecting wet-lab protocols to computational outputs, enhancing reproducibility and supporting automated workflow validation.
- Benchmarking with defined mock communities demonstrates workflow reliability across assembly methods while generating FAIR-compliant research objects suitable for community reuse and comparative studies.

## Introduction

Recent advances in high-throughput (omics) experimental methods have tremendously increased our ability to interrogate complex microbial communities and biological systems at the molecular level [1]. A FAIR end-to-end omics workflow produces three main types of metadata: source and sample material metadata, describing the experimental conditions under which raw omics data were obtained, software-centric operational metadata, describing details of software tools and parameter settings used, and omics product quality metadata describing intrinsic properties of the data products including accuracy, completeness, and consistency. Given the numerous methods developed for deriving information from raw data, each with its own strengths and weaknesses, rigorous tracking of the data provenance is essential.

Although machine-actionable minimal information models for metadata, designed to capture structured details of experimental research conditions, are currently being implemented [1], further extension of the FAIR principles to follow-up data processing workflows would greatly facilitate direct reuse of processed data, extracted information, and derived knowledge. The ISA metadata model [2] captures experiment metadata at three levels: ‘Investigation’, ‘Study’, and ‘Assay’ (Figure 1). The Investigation level provides conceptual and project administrative information and is mostly intended for human consumption. The Study and Assay levels outline the research design type (e.g., a time series design), the library source (e.g. metagenomic), the library strategy (e.g. WGS: Whole Genome Sequencing), and the technology platform used (e.g., OXFORD NANOPORE). These types of research metadata, play a role in directing the processing of raw data. Timely registration of this information in a machine-actionable format can streamline subsequent data processing and analysis, thereby enhancing overall efficiency.

**Figure 1.**
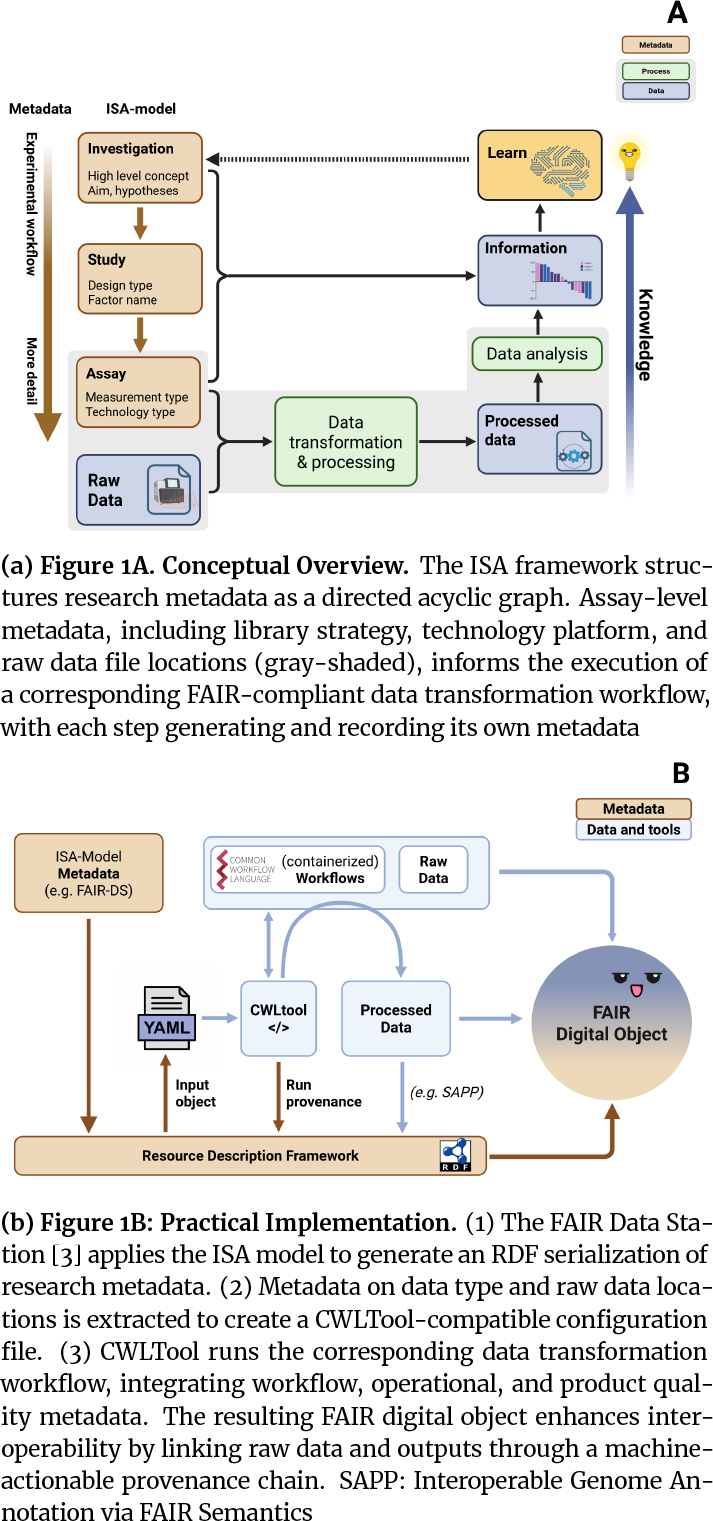
FAIR end-to-end workflows for ‘omics data analysis.

### Metadata managed data transformation workflows

Well-documented deterministic workflows are essential for reliably transforming raw outputs from omics experiments (e.g., raw DNA sequence libraries) into interpretable data products (e.g., sequence assemblies). A FAIR workflow should produce both product-related metadata, such as quality metrics (e.g., assembly completeness; see [4]), and process-related metadata that document the tools and parameters used.

Scientific computational workflows commonly use a stepwise approach with each step relying on a distinct software package. Given that each package has its own lifecycle and dependency requirements, a continuing challenge is finding those resources whose capabilities provide the best match with the requirements of the workflow. To address this ongoing issue, a common practice is to containerize each software package along with its dependencies, ensuring isolated and reproducible execution environments [5].

When each container is designed to address a single concern, it can be easily replaced with updated versions or with superior alternatives. This modularity also enables each workflow step to independently generate and report its own metadata. For instance, our current Metagenome-Assembled Genomes (MAG) reconstruction workflow (Figure 2) employs metaFlye [6] version 2.9.3, encapsulated within a Docker container sourced from BioContainers [7], for assembling Nanopore sequencing data. When only Illumina sequence data are available, metaSPAdes [8] version 3.15.5, is used. Documenting these specifications is essential to warrant the reproducibility of the resulting data products.

**Figure 2.**
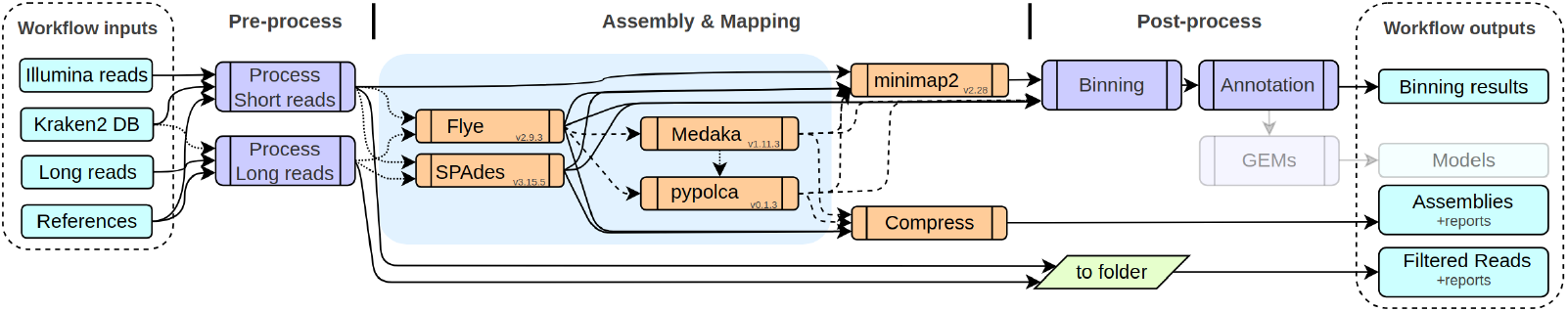
Schematic overview of a containerized modular workflow for MAG and GEM reconstruction from raw sequence reads. Workflow inputs and outputs are indicated blue. Orange steps represent command line tools encapsulated in a container. Two assembly tools (Flye, SPAdes) are integrated. When long reads are available, by default, the Flye assembly is used for binning. Nested workflows are in purple (collapsed, for brevity). Note, greyed out modules were not used in this study. CWL expressions for preparing output folders are in green. Workflow details are available at https://workflowhub.eu/workflows/367

Here, we demonstrate how interoperable experimental research metadata can be fused with follow-up FAIR4RS [9] computational processes. We show that such an integrated approach results in an expanded provenance chain, enhancing the traceability and FAIRness of the research objects obtained. We do this by evaluating the repeatability of a containerized modular workflow. The process involves the reconstruction of metagenome-assembled genomes (MAGs) using raw long- and short-reads from simulated microbial communities followed by FAIR-compliant comparative functional genome annotation using SAPP [10].

## Results

### Technical implementation

Metagenome assembly proceeds from raw reads through quality control, error correction, assembly, and polishing, followed by binning, MAG refinement, functional annotation and quality assessment. As described in Figure 1, a FAIR end-to-end metagenomics assembly workflow comprises two core components: a system to register experimental research metadata in an interoperable format and a follow-up data processing workflow capable of generating its own provenance. To enable FAIR registration of experimental metadata, we developed the FAIR Data Station (FAIR-DS), which implements the ISA data model in conjunction with the Genomic Standards Consortium’s minimum information standards to ensure machine-readable reporting of measurements and observations [3]. Originally designed to support metadata management for new experiments, FAIR-DS has since been extended and is used here to extract metadata from two BioProject studies registered at the European Nucleotide Archive (ENA).

Our data transformation workflows are defined using the Common Workflow Language (CWL, https://www.commonwl.org), an open standard for designing workflows that involve a sequence of command-line tools. Containerization of these tools supports workflow portability, parallelization and shareability. Tool encapsulation supports separation of concerns, promoting maintainability and repeatability. All our current workflows are hosted on WorkflowHub [11] (https://workflowhub.eu).

The workflows developed for MAG reconstruction, (Figure 2) (Table 1) implement mainstream and community-supported software tools. Each workflow starts with a rigorous quality control process: For Nanopore reads, NanoPlot [12] generates quality control plots and filtlong [13] trims and filters low-quality reads. Minimap2 [14] is used to filter reads from known contaminants. When Illumina reads are available, FastQC [15] provides metrics on per-base sequence quality, GC content, and adapter sequences. fastp [16] then performs quality trimming, adapter removal, and filtering of low-quality reads. BBMap [17] is used to filter reads from known contaminants. Kraken2 [18] is used for taxonomic classification.

**Table 1.** Overview of assembly-polishing configurations tested in this study. The deterministic (-d) mode uses the –deterministic flag in Flye, which enforces single-threading reducing certain sources of variability.

| Assembler (Mode) | Polishing Strategy |
| --- | --- |
| metaSPAdes (deterministic by default) | None |
| metaFlye (default) | Medaka |
| metaFlye (-d) | Medaka |
| metaFlye (-d) | PyPolCA |
| metaFlye (-d) | Medaka + PyPolCA |

By default, long-read assembly is performed with metaFlye, followed by consensus polishing with Illumina reads; for this step, we assessed both Medaka [19] andor PyPolCA [20]. QUAST [21] is used to assess assembly quality with metrics like N50, total length, number of contigs, and GC. During the binning stage multiple algorithms, MetaBAT2 [22],

MaxBin2 [23], and SemiBin2 [24] were applied to group contigs into MAGs. DAS Tool [25] integrates results from these binning algorithms to produce an optimized, non-redundant set of bins. Finally, CheckM [26] and BUSCO [27] assess the completeness and contamination of the binned MAGs, while GTDB-Tk [28] provides taxonomic classification for phylogenetic placement. In parallel, the CWLprov ontology, integrated within CWLtools [29], captures operational workflow metadata, recording each processing step along with input configurations, software versions, and their dependencies.

### Workflow performances and reproducibility assessment

To systematically evaluate the performances and reproducibility of the assembly workflow, we tested five assembly–polishing configurations.(Table 1). We assessed each configuration for performance, reproducibility, and its ability to handle different levels of data complexity using three simulated metagenomic datasets; ZYMO-even (CSI), ZYMO-log (CSII) [30] and BMOCK12 [31]. The bioproject numbers registered in ENA (Table 2) were used by the FAIR-DS to automatically retrieve metadata and data set locations of the three simulated metagenomic datasets in Turtle serialization and Excel format (Supplementary File S1 and S2). Standard SPARQL queries were applied to extract metadata essential for downstream processing; library source (METAGENOMIC), library strategy (WGS), sequencing platforms, (OXFORD_NANOPORE and ILLUMINA) (Table 2), and the link to the location of the corresponding raw sequence data files as input for the CWL tool definition configuration file (Supplementary File S4). For each assembly configuration, every dataset was processed in triplicate with operational metadata logged for each run. The resulting assemblies and bins were then compared to evaluate consistency across replicates.

**Table 2.** Mock microbial communities (meta) data used for benchmarking.

| Bioproject | Description |
| --- | --- |
| PRJEB29504 | **ZYMO (Even Distribution):** Mock microbial community standard, ultra-deep, Even: Eight bacterial species, even contribution based on genomic DNA, and two yeast species contributing 2% each; long-read nanopore & short-read Illumina available. |
| PRJEB29504 | **ZYMO (Log Distribution):** Mock microbial community standard, ultra-deep, eight bacterial and two yeast species, log distribution based on genomic DNA. Long read nanopore & short read Illumina available. |
| PRJNA496047 | **BMOCK12:** Mock microbial community for the benchmarking purposes. 12 isolate genomes from Actinobacteria, Proteobacteria and Bacteroidetes phyla; long read nanopore & short read Illumina. |
| **Zymo-LOG metadata RDF snippet in Turtle syntax. Essential (traversal) metadata in bold\*** |  |
| RDF Snippet | `<http://fairbydesign.nl/ontology/inv_fdp/ stu_ZYMO_PRJEB29504obs_ZYMO_EVEN_mocktest_cwl/ sam_Zymo-LOG_community_ONT/asy_ONT_LOG_ERR3152366> a jerm:Assay ; fair:alias "ena-RUN-UB-12-02-2019-16:02:39:937-3" ; fair:ena-base-count "16506755978" ; fair:ena-first-public "2019-02-12" ; fair:ena-last-update "2019-02-13" ; fair:ena-spot-count "3667480" ; fair:facility "Sequence facility is not available" ; fair:file_bytes "17147527495" ; fair:file_md5 "ffbb0daf168c08a6c4c9393481546bcc" ; fair:flowcell "unknown" ; fair:instrument_model "GridION" ; fair:library_selection "RANDOM" ; fair:library_source "METAGENOMIC" ; fair:library_strategy "WGS" ; fair:packageName "Nanopore" ; fair:platform "OXFORD_NANOPORE" ; fair:protocol "Protocol is not available" ; fair:sequencing_kit "unknown" ; schema:dataset <file://ftp.sra.ebi.ac.uk/vol1/fastq/ERR315/006/ERR3152366/ERR3152366.fastq.gz> ; schema:dateCreated "2019-02-12"; schema:description "GridION sequencing of ZYMOBIOMICS Microbial Community log distributed community"; schema:identifier "ONT_LOG_ERR3152366" .` |

\* The complete RDF output is provided in Supplementary File S1. For permitted values, see: https://ena-docs.readthedocs.io/en/latest/submit/reads/webin-cli.html

#### Workflow operational metadata

Workflow operational metadata confirms whether the workflow ran as expected. For this reason, output assessment should be preceded by a review of key operational metadata, including timestamps and the runtimes of individual tools. Table 3 presents operational metadata from two representative Flye BMOCK12 test runs, showing the impact of the –deterministic (-d) flag (which enforces single threading) on Flye’s runtime, on our current hardware; approximately 2.5 hours (-d) compared to 46 minutes with multi-threading enabled. A complete overview of all runs and the corresponding SPARQL query used to obtain these metadata are provided in Supplementary File S5.

**Table 3.** Operational metadata extracted from software tools used in the BMOCK12 Flye–Medaka assembly workflow. Two Flye assembler setting were compared: deterministic (A) and non-deterministic (B). Each setting was run in triplicate and for each software tool name and version and mean, maximum, and minimum (cumulative) execution times were extracted. Run times are reported in ISO 8601 format. Note that a single run may involve multiple uses of the same software tool.

| A: Deterministic mode |  |  |  |  |
| --- | --- | --- | --- | --- |
| Step | Tool Name | Version | Avg Run Time | Max/Min Run Time |
| Read Filtering and Classification | Fastqc | 0.12.1 | PT47M16S | PT47M52S / PT46M28S |
|  | Fastp | 0.23.4 | PT13M30S | PT13M59S / PT12M39S |
|  | BBmap | 39.06 | PT13M33S | PT17M / PT10M10S |
|  | Filtlong | 0.2.1 | PT17M22S | PT17M31S / PT17M14S |
|  | NanoPlot | 1.43.0 | PT7M54S | PT8M1S / PT7M47S |
| Assembly | Flye | 2.9.3 | PT2H36M1S | PT2H44M15S / PT2H25M59S |
|  | Medaka | 1.11.3 | PT10M10S | PT11M12S / PT9M29S |
|  | Quast | 5.2.0 | PT2M15S | PT2M36S / PT2M3S |
|  | BUSCO | 5.7.0 | PT8M6S | PT8M20S / PT7M49S |
| Annotation | Bakta | 1.9.4 | PT13M37S | PT13M41S / PT13M32S |
|  | Kofamscan | 1.3.0 | PT12M57S | PT13M18S / PT12M14S |
|  | Interproscan | 5.69-101.0 | PT29M54S | PT30M6S / PT29M47S |
|  | SAPP | 2.0 | PT23M20S | PT23M27S / PT23M11S |
| Binning | Maxbin2 | 2.2.7 | PT36S | PT39S / PT31S |
|  | Metabat2 | 2.15 | PT4M5S | PT4M5S / PT4M5S |
|  | Semibin2 | 2.0.2 | PT1M22S | PT1M27S / PT1M19S |
|  | Eukrep | 0.6.7 | PT33S | PT33S / PT33S |
|  | DASTool | 1.1.5 | PT2M5S | PT2M40S / PT1M37S |
| Avg Time of a Run | PT8H15M43S |  |  |  |
| B: Non-Deterministic mode |  |  |  |  |
| Step | Tool Name | Version | Avg Run Time | Max/Min Run Time |
| Read Filtering and Classification | Fastqc | 0.12.1 | PT54M40S | PT59M32S / PT52M9S |
|  | Fastp | 0.23.4 | PT16M2S | PT16M18S / PT15M29S |
|  | BBmap | 39.06 | PT14M7S | PT22M / PT9M43S |
|  | Filtlong | 0.2.1 | PT19M13S | PT21M41S / PT17M50S |
|  | NanoPlot | 1.42.0 | PT15M42S | PT16M27S / PT15M11S |
| Assembly | Flye | 2.9.3 | PT46M13S | PT46M20S / PT46M8S |
|  | Medaka | 1.11.3 | PT9M31S | PT11M38S / PT8M21S |
|  | Quast | 5.2.0 | PT2M3S | PT2M28S / PT1M43S |
|  | BUSCO | 5.7.0 | PT9M59S | PT12M32S / PT8M32S |
| Annotation | Bakta | 1.9.4 | PT16M50S | PT19M5S / PT15M6S |
|  | Kofamscan | 1.3.0 | PT20M24S | PT26M57S / PT13M33S |
|  | Interproscan | 5.69-101.0 | PT1H5M28S | PT1H30M22S / PT29M30S |
|  | SAPP | 2.0 | PT25M53S | PT28M5S / PT24M29S |
| Binning | Maxbin2 | 2.2.7 | PT41S | PT46S / PT38S |
|  | Metabat2 | 2.15 | PT4M15S | PT4M25S / PT4M9S |
|  | Semibin2 | 2.0.2 | PT1M46S | PT1M51S / PT1M37S |
|  | Eukrep | 0.6.7 | PT39S | PT42S / PT35S |
|  | DASTool | 1.1.5 | PT2M7S | PT2M55S / PT1M42S |
| Avg Time of a Run | PT7H43M18S |  |  |  |

\* Analysis where Illumina read data involved,
\*\* Analysis where ONT long read data involved

#### Assembly and binning quality metrics across datasets

Table 4 presents a comprehensive overview of assembly and binning performance across all five assembly configurations (Table 1), of the three mock community datasets, run in triplicate. The table serves as the central reference for comparing assembly contiguity, binning completeness, and reproducibility. Each dataset presents distinct challenges: ZYMO-EVEN offers uniform species distribution, ZYMO-LOG presents logarithmic abundance gradients spanning several orders of magnitude, while BMOCK12, composed of 12 bacterial species with varying abundances, provides a more complex and realistic benchmark for evaluating workflow robustness.

**Table 4.**
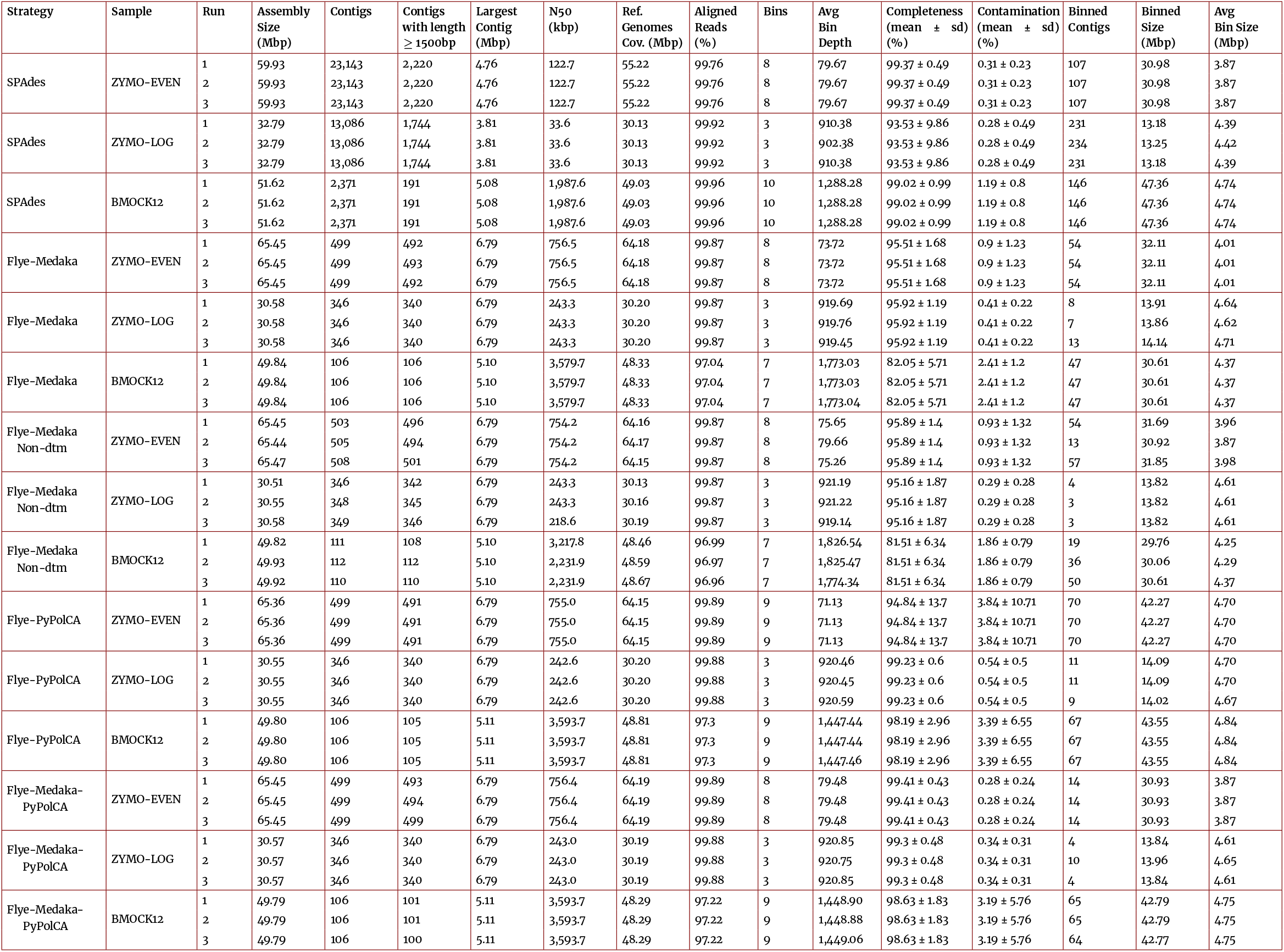
Summary of assembly and binning quality for different datasets using various assembly strategies.

For the ZYMO datasets, assembly contiguity metrics were notably better for long-read approaches, with both Flye-Medaka and Flye-PyPolCa producing a largest contig of 6.79 Mbp across both EVEN and LOG distributions. Overall the ZYMO-EVEN dataset, with its uniform species distribution, consistently yielded higher quality MAGs compared to ZYMO-LOG. For Flye-based assemblies, ZYMO-EVEN achieved 95.51% completeness with Medaka polishing, which improved to 99.41% with the addition of PyPolCA. In contrast, the logarithmic abundance distribution in ZYMO-LOG created challenges for binning algorithms, resulting in only 3 recovered bins compared to 8 bins from ZYMO-EVEN, reflecting the difficulty of reconstructing low-abundance species. The impact of setting the –deterministic (-d) flag is also evident: comparing Flye-Medaka (-d) with Flye-Medaka non-deterministic (-Non-dtm) revealed minimal differences in assembly metrics but between replicate runs a higher consistency in the number of contigs was obtained, e.g., 346, (deterministic) vs. 346-349 (-Non-dtm) for the ZYMO-LOG dataset. Note that Fly assembly outputs still can differ across runs, as multi-threading is not the only source of non-determinism in complex assembly graphs [32].

For the BMOCK12 dataset, long-read assembly using FlyePyPolCA produced highly contiguous assemblies with a largest contig of approximately 5.10 Mbp and an N50 of 3.58 Mbp, reflecting successful chromosome-level reconstruction. SPAdes short-read assembly yielded slightly longer total assembly sizes (51.62 Mbp vs. 49.80 Mbp) but with substantially more fragmentation; 2,371 contigs in total of vs. 106 for Flye. The Flye assemblies, while more contiguous, showed a lower MAG completeness, suggesting a trade-off between assembly contiguity and binning completeness. Notably, the polishing strategy significantly impacted final MAG quality: configurations incorporating PyPolCA (Flye-PyPolCA and Flye-Medaka-PyPolCA) achieved substantially higher completeness (98.19% and 98.63%, respectively) than Medaka alone, demonstrating PyPolCA’s effectiveness in using short-reads for error correction. However, these improvements came at the cost of slightly elevated contamination rates (3.39% and 3.19% vs. 2.41% for Flye-Medaka), likely reflecting increased sensitivity in detecting closely related strains or the presence of strain-level heterogeneity in the mock community.

Illumina read mapping rates were consistently high across all assembled datasets (≥96.96%), indicating strong concordance between long-read assemblies and short-read data. For BMOCK12, mapping rates ranged from 96.96-97.30% for Flye-based assemblies and 99.96% for SPAdes, confirming the accuracy of the assembled sequences. These mapping rates provide confidence in subsequent binning and polishing steps, as they demonstrate that the vast majority of short reads can be reliably aligned to the long-read assembly backbone.

The binning process captured a large proportion of the contigs, however there are differences. For example, in the BMOCK12 dataset, SPAdes assigned 146 of the 191 contigs larger than 1500 bp to 10 bins, capturing 91.17% of the assembled reads. In contrast, Fly-PyPolca assigned 67 of the 105 contigs larger than 1500 bp to 9 bins for the same dataset.

The species reconstruction heatmaps (Figure 4) further illustrated the consistency of species recovery across different assembly configurations and reveal important quality patterns related to community complexity. For the ZYMO-EVEN dataset, all assembly strategies successfully recovered the expected bacterial species, with most MAGs achieving a medium to high quality status according to MIMAG standards (completeness >90%, contamination <5%). The SPAdes and Flye-(Medaka)-PyPolCA configurations consistently produce high-quality MAGs for the major bacterial species. However, the two eukaryotic members (*Saccharomyces cerevisiae* and *Cryptococcus neoformans*) posed challenges for the standard prokaryotic binning algorithms [33], with variable recovery across configurations. In the ZYMO-LOG dataset, where species abundances span several orders of magnitude, only the high-abundance species were consistently recovered across all methods.

The BMOCK12 dataset, representing a more realistic microbial community scenario, presented the most complex pattern. The presence of two *Halomonas* strains (HL-93 and HL-4) in the mock community created challenges for the binning algorithms, resulting in hybrid bins harboring contigs from both strains (Table 4).

#### Average nucleotide identity analysis validates assembly accuracy

ANI analysis quantifies the species level accuracy of reconstructed MAGs by comparing them to their reference genomes. ANI values above 95% typically indicate species-level identity [34], while values approaching 99-100% suggest near-perfect genome reconstruction. The results demonstrate that all assembly strategies achieved high ANI values (>98%) for most recovered MAGs. ANI matrices for all three datasets are provided in Supplementary File S7, with ANI heatmaps shown in Supplementary Figures S1 (ZYMO-EVEN), S2 (ZYMO-LOG), and S3 (BMOCK12). Small variations in ANI values were observed across the different polishing methods, with the two-stage polishing approach typically resulting in marginally higher ANI values, reflecting the enhanced base-level accuracy obtained by integrating long-read and short-read consensus polishing.

ANI measures sequence similarity in shared regions but is insensitive to strain specific gene content differences. Consequently, the BMOCK12 hybrid *Halomonas* bin showed high ANI to both (HL-93 and HL-4) reference strains. This finding underscores a fundamental challenge in metagenomic binning: when closely related strains share >97-98% sequence identity [35], standard binning algorithms based on tetranucleotide frequencies and coverage patterns [22] may struggle to distinguish them cleanly.

### Functional annotation reveals strain-level diversity and chimeric assembly signatures

SAPP is an automated genome annotation platform that stores annotations and their provenance in a Linked Data format, enabling Semantic querying, integration, and mining [10]. Pfam domain based functional annotation of reconstructed MAGs provided insights into their metabolic potential and genomic composition. Individual MAGs exhibited highly conserved core metabolic functions typical of their respective taxonomic groups, including central carbon metabolism, amino acid biosynthesis, and energy conservation pathways (Supplementary File S9).

Domain-level analysis of *Halomonas* bins obtained from the BMOCK12 dataset suggested that this bin captured domains of both reference strains. We used Pfam domain presence/absence patterns to construct a Dollop parsimony tree (Figure 5) of *Halomonas* bins and the SPAdes bin appears to be the closest match to the reference strain. Quantitative analysis of the Pfam domain repertoires provided further insight into the functional resolution achieved by different assembly strategies. The two *Halomonas* reference genomes share a core domainome of 2,255 Pfam domains (HL-4: 2,306 domains; HL-93: 2,296 domains), representing 97.8% functional overlap between strains. The remaining 92 strain-specific domains (51 unique to HL-4, 41 unique to HL-93) served as diagnostic markers for assessing strain-level resolution in the assembled bins. Notably, the SPAdes assemblies recovered 63 of these 92 strain-specific domains (68.5%), demonstrating that the workflow captures not only the conserved core functional content but also a substantial fraction of the accessory domainome. Of the recovered strain-specific domains, 42 (66.7%) are associated with EC numbers, indicating enzymatic functions that could contribute to strain-specific metabolic capabilities.

Overall, bins derived from assemblies with lower contamination scores (Table 4) showed more coherent functional profiles, while those with higher contamination but better completeness exhibited evidence of foreign domain acquisition. The DAS Tool consensus binning approach generally selected bins that balanced both genomic completeness and functional coherence, as evidenced by bin identifiers showing preferential selection of MetaBAT2 or SemiBin2 results for different taxa, depending on which algorithm achieved better discrimination for that particular community member.

## Discussion

Metagenomics allows direct analysis of microbial communities from environmental samples, overcoming cultivation limitations and uncovering unculturable diversity. It reveals both community composition and functional capabilities, supporting the discovery of new genes and biological pathways. To improve reproducibility and FAIRness in metagenomic analyses, we designed a metadata-driven end-to-end framework that combines a FAIR experiment data collection with widely used sequence assembly, binning, and analysis tools, embedded within a FAIR-compliant containerized, end to end computational workflow. Its modular design incorporates metadata logging for workflow reproducibility, recording the tools, tool versions, and input parameters used during each run. This structure enables easy comparison tools and provides the flexibility to adapt the workflow to different input data types and analytical objectives. We validated the performances and robustness of the resultant workflows using triplicate runs of three mock communities with increasing complexity while its modularity allowed for a systematic assessment of various assembly-polishing configurations.

For the assembly step we compared SPAdes, a short-read assembler, with optional support for hybrid assembly using long reads with Flye, a genome assembler optimized for long-read data containing complex repeats. While the SPAdes assembler is deterministic by default the Fly algorithm is not. We studied the effects of its socalled deterministic mode (-d), which enforces single-threading and thus minimizes specific sources of variability. Beyond the anticipated increase in runtime, the results across all three mock datasets showed reduced variability in both the total number of contigs and assembly size between runs (Table 4). Overall, SPAdes consistently produced higher completeness scores across all datasets, with BMOCK12 achieving 99.02% completeness compared to 82.05% with Flye-Medaka. However, Flye-Medaka assemblies showed lower contamination rates (2.41% vs 1.19% for BMOCK12), suggesting a trade-off between completeness and contamination that researchers should consider when selecting assembly strategies.

Compared to short read data, long-read sequencing data typically have higher error rates and for polishing and further refinement of long read assemblies we compared Medaka, a polishing tool specifically designed for Oxford Nanopore sequencing data. [19] and (Py)PolCA a generic polishing tool for genome assemblies.that uses short-read sequencing data to increase consensus accuracy [20], [36]. We tested both tools separately and both proved to be essential for improving the quality of long-read assemblies. In addition, we tested a two-stage polishing approach (Flye-Medaka followed by PyPolCA) combining the potential benefits of nanopore-specific and short-read consensus corrections and noted a marginal increase in completeness and concommitant reduction of contamination with both tools in tandem ((Table 4).

During the data transformation process, two types of metadata are collected: product quality-related metadata (such as BUSCO scores as an estimate of genome completeness) and operational metadata (such as workflow runtime). We developed a set of SPARQL queries, implemented as Jupyter Notebooks, that can be used to generate customized reports for each metadata type. These notebooks are deposited in the GitLab code repository to ensure transparency and facilitate reuse. For product-based metadata, we used the Minimum Information about a Metagenome-Assembled Genome (MIMAG) standard [4] developed by the Genomic Standards Consortium (GSC) for reporting bacterial and archaeal genome sequences as a guideline. MIMAG extends the Minimum Information about Any (x) Sequence (MIxS) standard, which is also adopted by the FAIR-DS for FAIRifying raw input data. Comprehensive MIMAG/MIxS-compliant metadata reports for all MAGs are provided in Supplementary File S3. Metadata reports related to the computational transformation process are designed to monitor the details and performance of each workflow step compared to previous runs and runs with mock datasets. This approach helps to establish trust in the product-related output.

### MAGs as FAIR digital objects: Enabling provenance-based trust assessment

An important conceptual insight from our study concerns the inherent nature of metagenome-assembled genomes as consensus sequences. The observation of strain hybridization in the BMOCK12 *Halomonas* bins reveals a fundamental characteristic of metagenomic assembly: all MAGs represent some form of consensus genome, though the degree and nature of this consensus varies with the assembly strategy employed. This phenomenon warrants careful consideration when interpreting MAG-derived biological insights.

A central innovation of our approach is the conceptualization of MAGs not merely as sequence files, but as comprehensive FAIR digital objects that bundle genomic data with rich contextual metadata and complete provenance chains. Each MAG is accompanied by standardized quality metrics in accordance with the MIMAG standard [4], detailed operational metadata describing every computational step, and machine-readable provenance that links the final outputs to the original raw sequencing data and experimental design. This approach converts MAGs from opaque data products into transparent, self-documenting research objects that enable systematic trust assessment.

Despite the promising results, we acknowledge the limitations and challenges of our study. The binning of contigs from complex metagenomic samples remains a significant challenge, particularly for species with high strain-level diversity, low abundance, or large repeats. The long runtimes of some tools, such as InterProScan, can also be a bottleneck for high-throughput analyses.

### Potential implications

The machine-actionable nature of our metadata is particularly significant for enabling computational reasoning about data quality and fitness-for-purpose. By encoding all experiment metadata, assembly parameters, polishing strategies, binning algorithms, annotation and quality metrics in structured RDF format, we create datasets that can be interrogated programmatically to assess their suitability for specific downstream applications. For instance, a researcher or algorithm could automatically query the provenance graph to determine whether a MAG was polished with short reads (indicating higher base-level accuracy), whether closely related strains were present in the source community (indicating potential consensus effects), or whether the assembly was performed deterministically (enabling exact reproducibility). This level of transparency is essential for establishing trust in derived data products, especially as the volume of publicly available MAGs continues to grow exponentially.

Looking ahead, we see several exciting avenues for future research. A deeper evaluation of the GEMs produced from the MAGs using tools like memote [37] will be essential for validating their predictive accuracy. We also plan to further improve our RDF-based analysis tracking and its integration with RO-Crate to create even more comprehensive and interoperable research objects. The use of large language models and retrieval-augmented generation (RAG) to assist in the detection of candidate degrader strains and the assignment of taxonomic classifications is another promising area of research.

## Methods and Materials

### Study design and materials

We evaluated a metadata-managed, containerized workflow for metagenomic assembly, binning, and functional analysis using two defined mock community standards as input materials: BMOCK12 (BioProject PRJNA496047) and ZymoBIOMICS (BioProject PRJEB29504; Even and Log distributions). These communities provide ground-truth species composition and reference genomes, enabling controlled assessment of assembly and binning performance (Table 2; Figure 3). To assess the repeatability and robustness of our workflow, we performed triplicate runs for each mock community dataset using the full raw read sets across five distinct assembly configurations (Table 1). For the Flye assembler, we tested both deterministic and non-deterministic modes to evaluate algorithmic variability.

**Figure 3.**
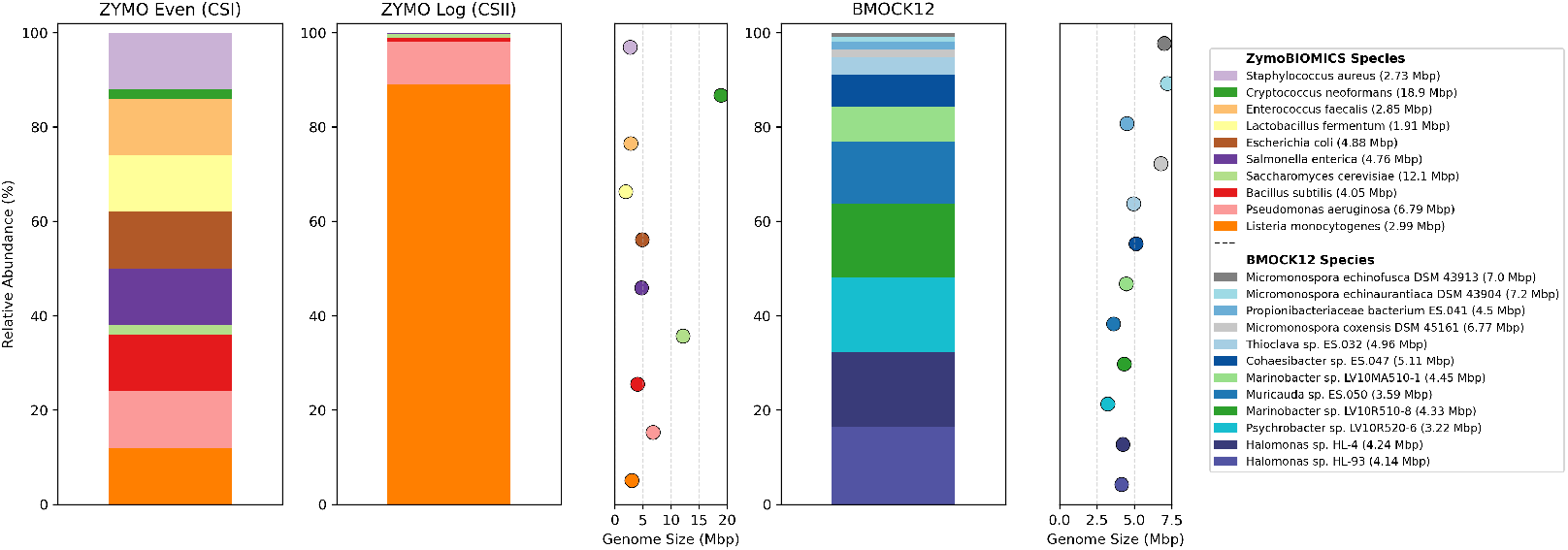
Composition, Relative Abundance, and Genome Size of Species in Mock Standards. ZymoBIOMICS species are ordered by their relative abundance in the CSII Mock standard. Scatter plots show corresponding genome sizes in Mbp.

### FAIR metadata capture and transformation

The experimental metadata for the mock communities were harmonized using the ISA (Investigation–Study–Assay) model and MIxS terms, as implemented in the FAIR Data Station (FAIR-DS) [3, 1]. The FAIR-DS generates an RDF/Turtle serialization of the metadata, where key traversal fields required to drive the computational workflow (e.g., library_source, library_strategy, platform, and raw data URIs) are explicitly defined (Table 2). This structured metadata is then automatically transformed into a CWLTool-compatible configuration file (in YAML format), which parameterizes and initiates the workflow run (Figure 1). This process creates a seamless, machine-actionable provenance chain that links the initial research objects, workflow definitions, parameter sets, and the final outputs.

### Workflow implementation and containerization

Our workflows are authored in the Common Workflow Language (CWL; https://www.commonwl.org) and executed using CWLTool. We adhere to the single-containment principle, where each command-line tool is encapsulated in its own container to isolate dependencies and enhance reproducibility [5, 11]. The container images are sourced from community registries like BioContainers and are referenced by immutable tags in the CWL tool descriptions. The complete workflow is publicly available on WorkflowHub (https://workflowhub.eu/workflows/367), and a summary of the software components and their versions, as captured at runtime, is provided in Table 3.

### Read quality control, filtering, and classification

For ONT reads, NanoPlot [12] was used to generate QC diagnostics, and Filtlong [13] was employed for trimming and filtering low-quality reads. If contaminant reference genomes were available, Minimap2 [14] was used to remove potential contaminant reads. For Illumina reads, FastQC [15] assessed quality metrics, while fastp [16] performed adapter and quality trimming. BBMap [17] was used for contaminant filtering. When applicable, Kraken2 [18] was used for the taxonomic profiling of reads. The exact versions and invocation parameters for all tools are recorded in the CWLProv provenance logs and are summarized in Table 3.

### Assembly and polishing

The assembly was performed using metaFlye [6] for long-read inputs and metaSPAdes [8] for short-read-only inputs, following the five distinct configurations outlined in Table 1. The long-read assemblies were polished with Medaka [19] and/or further corrected with PyPolCA [20] using the corresponding Illumina reads, depending on the specific configuration. The quality of the assemblies was assessed using QUAST [21], which reports key metrics such as N50, largest contig, total length, GC content, and the number of misassemblies. To investigate algorithmic determinism, Flye was executed both with settings that enforce determinism and in its default non-deterministic mode (FlyeMedaka-Nondtm configuration). The operational differences between these runs were captured by CWLProv and are summarized in Table 3.

### Binning and bin-set integration

To leverage the complementary heuristics of different algorithms, contigs were binned using multiple tools: MetaBAT2 [22], MaxBin2 [23], and SemiBin2 [24]. DAS Tool [25] was then used to integrate the candidate bins into a non-redundant, consensus bin set. Where relevant, EukRep [38] was applied to flag eukaryotic contigs prior to the binning process. Read-mapping statistics for both the assemblies and the bins were computed to estimate coverage and support downstream quality control.

### MAG quality assessment and taxonomic assignment

The quality of the Metagenome-Assembled Genomes (MAGs) was evaluated using CheckM and BUSCO [27] to quantify their completeness and contamination. The taxonomic classification of the MAGs was performed with GTDB-Tk [28]. We report the quality categories in accordance with the MIMAG standards [4] for bacterial and archaeal MAGs (Figure 4a and Figure 4b). When applicable, eukaryotic bins were handled separately from the bacterial and archaeal MAGs.

**Figure 4.**
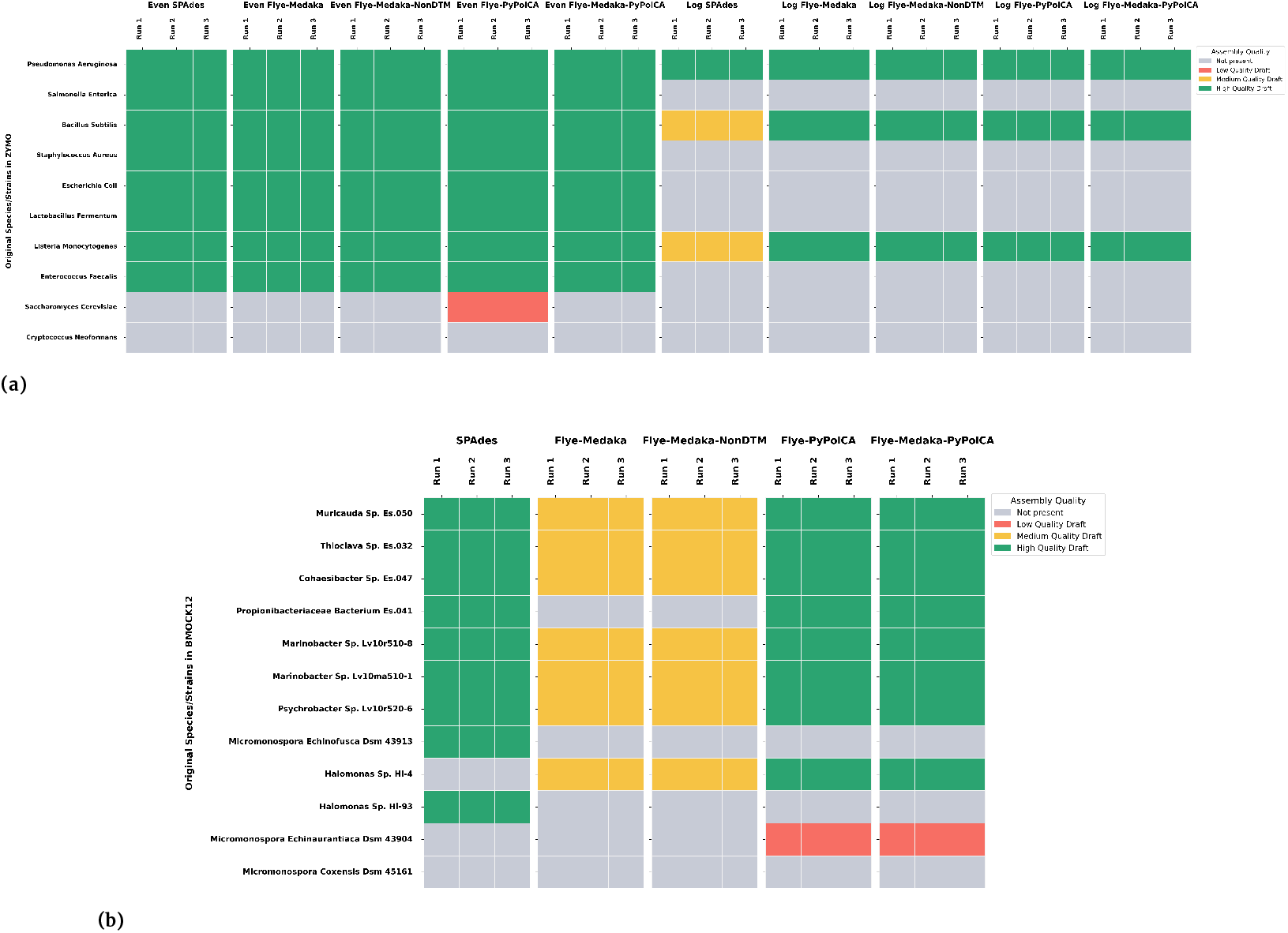
Heatmap of Reconstruction Presence and Assembly Quality of Species/Strains in Mock Community. (a) ZYMO Mock Community incl. Even and Log distribution. (b) BMOCK12 Mock Community. Quality categories are based on completeness, contamination, and rRNA/tRNA statistics as stated in MIMAG standards [4].

### Functional annotation

Gene prediction and functional annotation were performed using a combination of Bakta [39], KOfamScan [40], and InterProScan [41]. The annotations were then integrated in a FAIR-compliant manner with SAPP for interoperable functional semantics. The resulting domain-level profiles were used for comparative analyses, including clustering and the construction of phylogenetic trees (e.g., Dollop parsimony; Figure 5).

**Figure 5.**
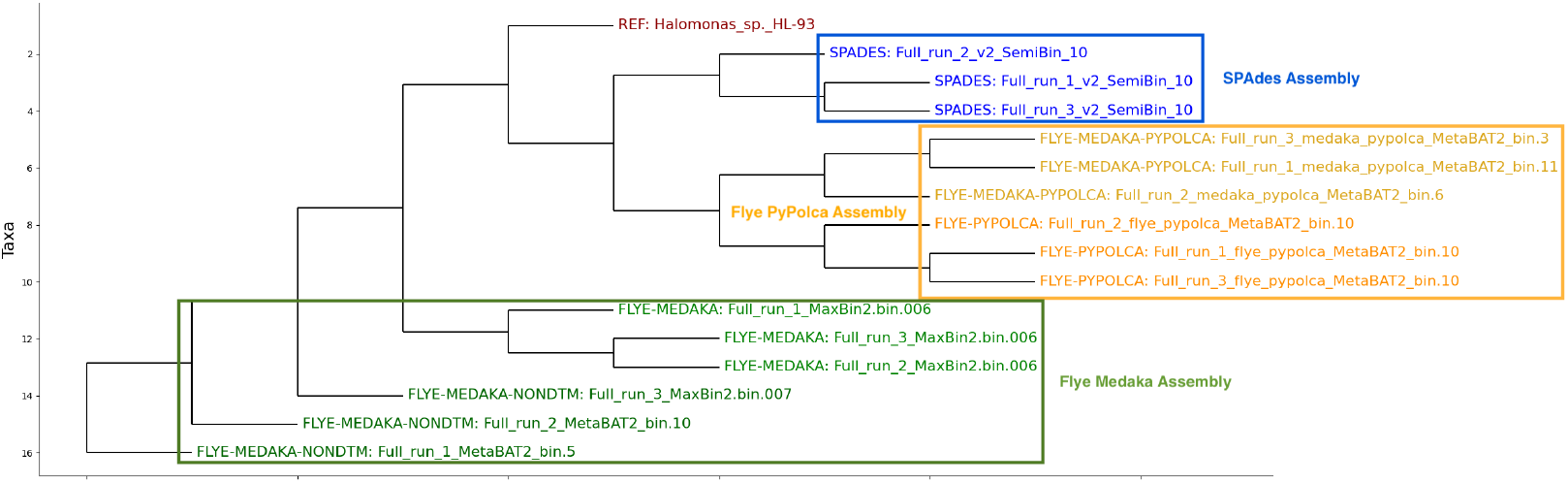
Phylogenetic analysis of *Halomonas* strains based on functional domain profiles. Dollop parsimony tree constructed from Pfam domain presence/absence binary matrix for *Halomonas* bins (HL-93) recovered from the BMOCK12 dataset. The *Halomonas* clade is positioned at the top. Some bins display evidence of chimeric composition, containing domains from both strains, consistent with the strain hybridization effect observed in quality metrics and ANI analysis.

### Provenance capture and FAIR packaging

During each workflow execution, CWLProv [29] recorded all inputs, outputs, software tools (including their versions and container hashes), parameter settings, start and end timestamps, and file-level derivations in W3C PROV-compatible RDF [42]. These provenance records were stored in GraphDB alongside functional annotation data (GBOL model; Supplementary Figure S4) for integrated querying and analysis. Each run was then packaged as a Research Object (RO-Crate) [43], which aggregates the primary outputs and the provenance metadata to ensure long-term reuse. The operational metadata and product-quality metrics were queried via SPARQL to generate run summaries (Table 3) and MIMAG-oriented reports. Complete RDF exports of both provenance and functional annotation data are provided as Supplementary Files S8 and S9, with the formal GBOL schema definition in Supplementary Files S10–S11, enabling researchers to load these data into their own RDF triple stores for custom queries and analysis.

### Reproducibility experiments

To quantify the repeatability of our workflow, each configuration (i.e., each dataset, assembler/polisher variant, and Flye mode) was executed in triplicate under identical conditions. We compared the results based on several key metrics: (i) assembly metrics (e.g., largest contig, N50), (ii) mapping rates of Illumina reads to ONT-assembled contigs, (iii) MAG counts and quality (completeness/contamination), and (iv) the Average Nucleotide Identity (ANI) of the bins to their reference genomes. For both the BMOCK12 and ZYMO Log datasets, we analyzed the full datasets.

### Computational environment

The workflows were executed on Linux x86_64 systems with Docker-compatible container runtimes. The resource usage and timing for each tool were recorded by CWLProv and are summarized in Table 3. The use of CWL and containerized tools ensures the portability of the workflow across different workstations and clusters. The orchestration of the workflow was performed by CWLTool, without reliance on any site-specific schedulers (e.g., Slurm, SGE).

### Statistical analysis and visualization

The post-processing of the workflow outputs and provenance data was performed using Python-based notebooks with the pandas, matplotlib, and seaborn libraries [44, 45, 46]. The ANI matrices were computed with FastANI [47] (via utility wrappers) and visualized as heatmaps. Raw ANI matrices for all datasets are provided in Supplementary File S7. Where appropriate, summary statistics, including the mean, minimum, and maximum values across the triplicate executions for a given configuration, are reported (Table 3).

### Software, parameters, and availability

All software tools are referenced in the workflow with versioned container images. The run-time, versions and parameters used are captured in CWLProv and are included in each RO-Crate. The MAG reconstruction workflow, including the CWL tool and workflow definitions, is publicly available on WorkflowHub (https://workflowhub.eu/workflows/367). Example configuration files derived from the FAIR-DS metadata are provided in Supplementary File S4. SPARQL query templates used to generate operational and product-quality reports are provided in Supplementary File S5. Complete operational metadata including runtime statistics for all workflow runs are provided in Supplementary File S6.

## Availability of source code and requirements

Lists the following:

- Project name: (Hybrid) Metagenomics workflow
- Project home page: https://workflowhub.eu/workflows/367
- Operating system(s): Platform independent
- Programming language: Python, Java, shell, R
- Other requirements: docker, cwltool, (kubernetes)
- License: APACHE 2.0

## Data availability

The data sets supporting the results of this article are available in the following repositories:

- Test datasets: The mock community datasets (BMOCK12 and ZYMO) used for validation are available from their original publications [31] and [30].
- Supplementary data files: The following supplementary files are deposited in the Zenodo repository [DOI: 10.5281/zenodo.17990145] [48] and are also included with this article:
  – **Supplementary File S1:** FAIR-DS experimental metadata in RDF/Turtle format, including ISA model structure and MIxS-compliant metadata for all mock communities.
  – **Supplementary File S2:** FAIR-DS experimental metadata in Excel format for human-readable access.
  – **Supplementary File S3:** MIMAG/MIxS-compliant metadata reports for all MAGs, including completeness, contamination, and taxonomic classification.
  – **Supplementary File S4:** CWL tool definition configuration files (YAML format) for all workflow runs.
  – **Supplementary File S5:** SPARQL query templates for extracting operational and quality metrics from GraphDB.
  – **Supplementary File S6:** Complete operational metadata for all workflow runs, including runtime statistics and tool execution times.
  – **Supplementary File S7:** Raw ANI matrices (pairwise values) for all three datasets.
  – **Supplementary File S8:** Complete workflow provenance data in RDF/Turtle format (PROV-O/CWLProv compliant).
  – **Supplementary File S9:** Functional annotation data in RDF/Turtle format (GBOL ontology).
  – **Supplementary File S10:** GBOL data model schema in Mermaid format.
  – **Supplementary File S11:** GBOL data model schema in ShEx (Shape Expressions) format.
  – **Supplementary Figures S1–S3:** ANI heatmaps for ZYMO-EVEN, ZYMO-LOG, and BMOCK12 datasets.
  – **Supplementary Figure S4:** GBOL schema class diagram showing the structure of functional annotation data. The RDF datasets (S8 and S9) can be loaded into any RDF-compatible triple store and queried using standard SPARQL tools. Example SPARQL queries for extracting specific information are provided in Supplementary File S5. The RDF data use standard ontologies (PROV-O [42], CWLProv [29], GBOL), ensuring interoperability and enabling integration with other FAIR-compliant datasets. The complete GBOL data model schema is provided in Supplementary File S10-S11 (Mermaid diagram - ShEx format) and visualized in Supplementary Figure S4.
- Workflow code and analysis notebooks: The workflow source code and Jupyter notebooks used for data analysis, figure generation, and table preparation are available on GitLab at https://git.wur.nl/unlock/projects/FAIRwf4MicrobialCommunity.
- Workflows: The workflow definitions are available on WorkflowHub [49] and their source code is hosted on GitLab at https://gitlab.com/m-unlock/cwl.

## Author’s Contributions

**CK**: Conceptualization, Data Curation, Formal Analysis, Investigation (benchmarking and data analysis), Methodology (workflow design & development), Software, Validation, Visualization, Writing & Review. **JJK**: Methodology (FAIR Data Station), Software, Supervision, Writing & Review. **BN**: Methodology (CWL pipelines and FAIR Data Station), Software. **WTS**: Investigation (GEM analysis), Writing & Review. **PJS**: Conceptualization, Funding Acquisition, Project Administration, Resources, Supervision, Writing & Review.

